# immuneSIM: tunable multi-feature simulation of B- and T-cell receptor repertoires for immunoinformatics benchmarking

**DOI:** 10.1101/759795

**Authors:** Cédric R. Weber, Rahmad Akbar, Alexander Yermanos, Milena Pavlović, Igor Snapkov, Geir Kjetil Sandve, Sai T. Reddy, Victor Greiff

**Affiliations:** Department of Biosystems Science and Engineering, ETH Zürich, 4058 Basel, Switzerland; Department of Immunology, University of Oslo, 0372 Oslo, Norway; Department of Informatics, University of Oslo, 0373 Oslo, Norway

## Abstract

**Summary:** B- and T-cell receptor repertoires of the adaptive immune system have become a key target for diagnostics and therapeutics research. Consequently, there is a rapidly growing number of bioinformatics tools for immune repertoire analysis. Benchmarking of such tools is crucial for ensuring reproducible and generalizable computational analyses. Currently, however, it remains challenging to create standardized ground truth immune receptor repertoires for immunoinformatics tool benchmarking. Therefore, we developed immuneSIM, an R package that allows the simulation of native-like and aberrant synthetic full length variable region immune receptor sequences. ImmuneSIM enables the tuning of the immune receptor features: (i) species and chain type (BCR, TCR, single, paired), (ii) germline gene usage, (iii) occurrence of insertions and deletions, (iv) clonal abundance, (v) somatic hypermutation, and (vi) sequence motifs. Each simulated sequence is annotated by the complete set of simulation events that contributed to its in silico generation. immuneSIM permits the benchmarking of key computational tools for immune receptor analysis such as germline gene annotation, diversity and overlap estimation, sequence similarity, network architecture, clustering analysis, and machine learning methods for motif detection.

**Availability:** The package is available via https://github.com/GreiffLab/immuneSIM and will also be available at CRAN (submitted). The documentation is hosted at https://immuneSIM.readthedocs.io.

## 1 Introduction

Targeted deep sequencing of adaptive immune receptor repertoires (AIRR-seq data, (Breden *et al.*, 2017)) has become a key resource for immunodiagnostics and immunotherapeutics research. Consequently, there exists a rapidly growing number of immune receptor informatics tools for germline gene annotation, diversity and overlap estimation, network architecture (sequence similarity), and machine learning analysis (Miho *et al.*, 2018; Greiff *et al.*, 2015; Yaari and Kleinstein, 2015; Brown *et al.*, 2019). In order to benchmark and assess the performance of these tools (Olson *et al.*, 2019), synthetic ground truth immune receptor datasets with complete information on all repertoire feature dimensions investigated or used in these tools (e.g., germline gene usage, insertion and deletions, and clonal abundance, Fig. 1) are required (Bashford-Rogers *et al.*, 2013; Yaari and Kleinstein, 2015; Madi *et al.*, 2017; Greiff, Weber, *et al.*, 2017; Miho *et al.*, 2017, 2018; Brown *et al.*, 2019). Therefore, there is a need for a computational framework that enables the simulation of native-like immune receptor repertoires and allows for the tracing of all immunologically-relevant simulation parameters. To address this gap in the landscape of immune receptor simulation tools (Safonova *et al.*, 2015; Marcou *et al.*, 2017; Yermanos *et al.*, 2017), we here present the immuneSIM R package, which allows the simulation of human and mouse BCR and TCR repertoires (single-chain and paired full length variable regions) with traceable simulation-event level annotation for each of the simulated sequences. The user has full control over the following immunological features: V-, D-, J-germline gene set [germline gene sets of species other than human or mouse or additional newly discovered germline genes may be added by the user] and usage frequencies, occurrence of insertions and deletions, clonal sequence abundance and somatic hypermutation. Post-sequence simulation, the generated immune receptor sequences may be further altered by the addition of custom sequence motifs, synonymous codon replacement as well as the modification of the sequence similarity architecture (Fig. 1). We validated that immuneSIM can generate immune repertoires that are similar to experimental repertoires (native-like) by evaluating a range of repertoire similarity measures. immuneSIM can also generate aberrant immune receptor repertoires to replicate a broad range of experimental, immunological or disease settings (Brown *et al.*, 2019; Arora *et al.*, 2019) (Supplementary Figs 2–7).

**Fig. 1.**
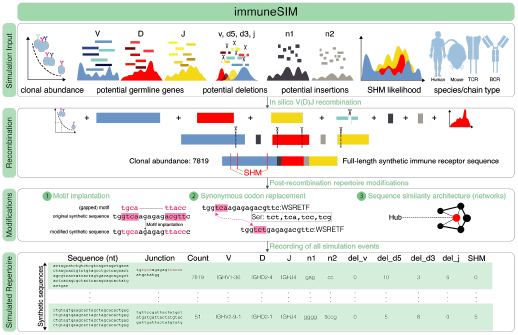
immuneSIM simulates fully (single, paired) annotated immune repertoires that are either highly similar (native-like) or deviating (aberrant, see main text for definition) from experimental immune repertoires. All major immune repertoire features such as clonal abundance, germline genes, deletions and insertions, and somatic hypermutation are tunable. Post in silico recombination, the immuneSIM-generated immune receptor repertoires may be further modified by (1) implantation of motifs, (2) codon replacement and (3) change of sequence similarity architecture.

## 2 Package Description

immuneSIM enables the simulation of native-like and aberrant repertoires for the benchmarking of immunoinformatics tools. In order to simulate immune receptor repertoires that mimic native-like repertoires, immuneSIM contains reference experimental data from human and mouse studies (Greiff, Menzel, *et al.*, 2017; Madi *et al.*, 2014; DeWitt *et al.*, 2016; Emerson *et al.*, 2017, Supplementary Table 1). ImmuneSIM allows for further customization by permitting the inclusion of alternative experimental or user-created reference datasets as well as the simulation of *aberrant* repertoires with feature distributions different from those observed in the input experimental parameters provided by the immuneSIM package. The in silico recombination process (Fig. 1, Supplementary Fig. 1) starts by sampling V, D and J genes according to a given frequency distribution (possibly sampled from input datasets), followed by the simulation of deletion events for the V and D genes. To increase the probability of providing the user with in-frame junctional regions, the J gene deletion length is chosen in such a way that the J-gene anchor (i.e., the nucleotide pattern that marks the J region of the CDR3) (Giudicelli *et al.*, 2004; Giudicelli and Lefranc, 2011) remains in-frame. Likewise, the n1 (5’ of D-gene) and n2 (3’ of D-gene) insertion sequences are sampled from a subset of observed insertion sequences to ensure the maximal probability of generating an in-frame sequence. Following the assembly of the V, n1, D, n2 and J fragments into a full V(D)J sequence, a clone abundance is assigned to it, and somatic hypermutation (for B-cell receptors only) based on the R package AbSim (Yermanos *et al.*, 2017) may be applied. Depending on the V,(D),J genes sampled, it is possible that the simulated sequences contain stop codons; any such unproductive sequences are automatically discarded. immuneSIM continues to simulate recombined sequences until the user-defined number of sequences has been reached. To reach very large numbers of diverse BCR and TCR repertoires, simulations may be parallelized (Supplementary Figs. 8, 9). Each immuneSIM-generated sequence is annotated with all simulation events that led to its generation including species (human, mouse), chain type (single, paired; TCRβ/TCRα, IgH/Igκ, Igλ), clonal abundance, V,(D),J germline gene usage, information on deletions, insertions (n1, n2) and junctional recombination (complementarity determining region 3, CDR3). Additionally, modifications such as synonymous codon replacement (relevant for testing the analytical performance of nucleotide and amino acid sequence-based methods) and alteration of repertoire sequence similarity architecture (relevant for testing graph-based tools) can be performed (Fig. 1). Further, a flexible motif implantation function allows the user to simulate specific sequence motifs through the controlled insertion of short sequence motifs of various complexities (k-mers of various sequence lengths and diversity at specified frequencies). Sequence motifs have been previously shown to be implicated in the prediction of public and private clones (Greiff, Weber, *et al.*, 2017) as well as antigen binding and disease course (Dash *et al.*, 2017; Glanville *et al.*, 2017; Ostmeyer *et al.*, 2019).

## 3 Validation: Simulating native-like and aberrant immune receptor repertoires

To validate the similarity between simulated and experimental repertoires, we simulated repertoires using standard settings based on experimental data as well as aberrant repertoires with parameters introducing noise in various dimensions (mouse, human, IGH and human TRB, see Supplementary Table 2). We first compared simulated B- and T-cell repertoires to experimental datasets in terms of CDR3 length distribution, VDJ usage, positional amino acid frequencies and k-mer co-occurrence (gapped subsequence structures) (Supplementary Figures 2–5). Briefly, we found that murine IgH repertoires simulated with the standard parameters replicate V, D, and J gene frequencies between input and output (r_Spearman_ ≥ 0.985), whereas aberrant simulated BCR and TCR repertoires showed larger deviations (r_Spearman_ ≥ 0.8). Similarly, the amino acid frequency distribution differed only slightly compared to the naïve repertoire when using standard parameters (mmse across positions = 0.000486) in contrast to aberrant repertoires that were more distant (mmse across positions = 0.001659). Finally, gapped-k-mer subsequence usage correlated highly between standard simulations and experimental repertoires (r_Spearman_ = 0.86, at k = 3, m ≤ 3, where k is the k-mer amino acid length and m is the number of amino acid gaps) while aberrant repertoires showed more distinct gapped k-mer patterns (r_Spearman_ = 0.74). To further substantiate the congruence of experimental and immuneSIM generated repertoires, we determined the extent to which the internal annotation of simulated repertoires overlapped with IMGT’s HighV-Quest (Alamyar *et al.*, 2012), a commonly-used annotation tool (Supplementary Figs. 6, 7). We found up to 99% of simulated sequences were annotated as ‘productive’ and ‘in-frame’ by IMGT HighV-Quest. Among these sequences, 94% of the time the junction identified by immuneSIM was found to be identical to that of IMGT. The V and J annotation overlapped in >97% of simulated sequences, while D annotations, a generally more difficult problem due to deletions and insertions (Bolotin *et al.*, 2015; Safonova and Pevzner, 2019), showed an overlap of ~60%. Taken together, these results support the notion that immuneSIM repertoires are nearly indistinguishable from experimental repertoires with respect to major statistical descriptors and thus can serve as a reliable basis for benchmarking immunoinformatics tools. Finally, immuneSIM may serve for tool stress-testing analysis, for example benchmarking machine learning methods (Emerson *et al.*, 2017; Greiff, Weber, *et al.*, 2017), using implanted sequence motifs at various frequencies and complexities. The versatility and flexibility of immuneSIM (BCR, TCR, single-chain, paired chain, multiple species, user-defined inputs of key immunological parameters) will be helpful for benchmarking the reliability and accuracy of informatics tools used to analyze immune repertoire deep sequencing data.

## Supplementary Data

### Detailed Methods

A detailed explanation of each function of the immuneSIM R package is provided in the online documentation https://immuneSIM.readthedocs.io. Implementation details and procedures relevant to the understanding of the results (Supplementary Figures 2–9) presented in the Supplementary Data are described in more detail below.

### Sampling of germline V, D, J genes

V, D and J genes are sampled based on V, D, J usage frequencies contained in the provided `list_germline_genes_allele_01` R list object contained in the package. The list contains IMGT germline gene sequences (Alamyar *et al.*, 2012) for human and murine immunoglobulin heavy and light and T-cell receptor beta and alpha chains as well as frequency distributions for each subset. In the case of immunoglobulin heavy and beta chain repertoires, the germline gene frequencies were obtained from published datasets (Greiff, Menzel, *et al.*, 2017; Madi *et al.*, 2017; DeWitt *et al.*, 2016; Emerson *et al.*, 2017), while uniform distributions were used in the case of light and beta chain repertoires due to a lack of fitting datasets (see Supplementary Table 1). The user is free to modify these frequencies or extend the `list_germline_genes_allele_01` to introduce different experimental frequencies or synthetic germline sequences. Since different V, D, J combinations have an unequal likelihood to form a productive in-frame sequence, a purely input-frequency-based sampling can lead to skewed V, D, J usage in simulated repertoires. Therefore, the `update_vdj_freqs` parameter allows for a re-adjustment of the initial VDJ frequencies after a defined threshold (default: 50% of the sequences) in order to generate sequences with a VDJ distribution that closely mirrors input frequencies.

### Sampling of deletions and insertions

The deletion lengths are sampled from a provided R dataframe `insertions_and_deletion_lengths_df` which contains deletion lengths and insertion sequences pooled across all samples of a previously published mouse BCR study (Greiff, Menzel, *et al.*, 2017). The immuneSIM R package contains a subset of 500’000 entries, while the full dataset (11 ‘363’603 entries) is available through GitHub (https://github.com/GreiffLab/immuneSIM) and can be loaded using the immuneSIM function `load_insdel_data ()`. Each row in this reference dataset contains an experimentally observed insertion and deletion event. The provided data does not contain information on correlations between insertion and deletions and specific germline genes in order to enable a wider variety of possible recombinations of insertions, deletions and germline genes. While immuneSIM does not provide a species or receptor chain specific reference dataset, the 11 ‘363’603 data points for n1, n2 insertions and V, D, J deletions cover a wide range of all theoretically possible insertion and deletion patterns. Analogously to the reference for the sampling of the V, D and J germline genes, the user has the option to modify or replace this reference file. The sampling of insertion and deletion starts with the sampling of the deletion lengths for the 3’ end of the V gene as well as on 5’ and 3’ end of the D gene from `insertions_and_deletion_lengths_df`. The deletion at the 5’ J region is applied after a valid anchor point is identified – J-TRP and J-PHE (represented by “`tgg`” and “`ttt`”, “`ttc`”, respectively) (Lefranc, 2011). Specifically, the deletion is applied to the pre-anchor subsequence of the chosen J gene. For each deletion length, an additional checkpoint ensures that they are not longer than the sequence they are applied to. After deletion lengths have been sampled and applied, the insertions n1 and n2 are sampled from a subset of `insertions_and_deletion_lengths_df` which is restricted to insertions of lengths that will result in an in-frame CDR3 (complementarity determining region 3).

### Identifying the CDR3 junctional region

The junctional region of the immune receptor sequence, at which the V, D and J genes are brought together, forms the CDR3. During the simulation of each sequence, the CDR3 is determined based on the V-gene cysteine anchor (represented by“`tgt`”, “`tgc`”) and the J-gene anchors J-TRP, J-PHE (Lefranc, 2011). This CDR3 is translated into amino acids and matched to the full sequence. If the sequence is productive (in-frame, no stop codons) an additional check is performed to determine whether the determined J-gene anchor is correct by looking for additional occurrences of out-of-frame patterns (as identified via High-VQuest analysis of simulated sequences) further downstream of the sequence. Following this, a further check ensures that the amino acid length of the CDR3 detected is not above or below a user-set threshold.

### Motif implantation

Since sequence motifs have been previously shown to be predictive of public and private clones (Greiff, Weber, *et al.*, 2017) as well as antigen binding and disease course (Dash *et al.*, 2017; Glanville *et al.*, 2017; Ostmeyer *et al.*, 2019), controlled and recoverable implantation of motifs is essential for the benchmarking of machine learning methods. immuneSIM enables the user to implant user-defined or randomly generated k-mer motifs at various frequencies in the nucleotide and amino acid CDR3 sequences. Additionally, there is an option to choose between positional and position-agnostic implantation. The immuneSIM function `motif_implantation` takes the user input consisting of an AIRR-compliant repertoire (Rubelt *et al.*, 2017), a list defining the motifs, their frequencies and a position parameter and implants the motif in the CDR3 of the nucleotide and amino acid sequences of the repertoire. At the implantation position, the existing nucleotides and amino acids are replaced, thus conserving the length of the modified CDR3.

### Synonymous codon replacement

In order to create repertoires that have 100% amino acid sequence identity but differ in their nucleotide composition (relevant for testing the predictive power of nucleotide and amino acid sequence-based methods), immuneSIM provides the function `codon_replacement`, which allows for the substitution of nucleotide codons. This can be relevant for testing predictive performance of methods for which both amino acid-based features (such as amino acid frequency distributions) as well as nucleotide-based features (e.g., gapped-k-mer occurrence) can be used as input. Specifically, an example of such a use case would be the prediction of public and private clones using Support Vector Machines (Greiff, Weber, *et al.*, 2017). The function takes as input (i) an AIRR-compliant repertoire (Rubelt *et al.*, 2017), (ii) the sequences that should be modified (“AA”, “nt” or “both”), (iii) a list containing the user-defined rules for replacement (i.e. the codons that should be replaced and their replacements) and (iv) a probability with which a sequence in the repertoire should be skipped in the replacement process. Based on the probability defined in (iv), each sequence is either marked as a target sequence (codon replacement is applied) or kept as is (no codon replacement occurs). The to-be-replaced codons are identified in each target sequence and all of them are replaced according to the rules specified in the user-provided list. While the main intended use case for the `codon_replacement` function is for synonymous replacement, it supports any kind of nucleotide replacement and can, therefore, also be used for nonsynonymous codon replacements that leads to a modification of the amino acid sequence.

### Architecture modification

The repertoire architecture as defined by the properties of the similarity network of its amino acid CDR3 sequences has been shown to be a reproducible and robust feature across B- and T-cell repertoires (Madi *et al.*, 2017; Miho *et al.*, 2019). The immuneSIM function `hub_seqs_exclusion` enables the modification of the similarity architecture through the deletion of the top X (percentage, defined by user) hub sequences (i.e. sequences of connective importance in the network). This is achieved by calculating and analyzing the CDR3 similarity network based on the Levenshtein distance (edit distance = 1) using the R packages igraph (Csardi and Nepusz, 2006) and stringdist (Loo, 2014). The top hub sequences are identified based on their hub score and excluded.

### Simulation of somatic hypermutation

The simulation of somatic hypermutation (SHM) is performed using the previously published AbSim R package (Yermanos *et al.*, 2017). After the sampling of the V, D, J genes and insertions and deletions, SHM is performed based on the user-defined mode and SHM probability (for details see package documentation). Each SHM event is recorded in a character string that is output in the repertoire dataframe. immuneSIM additionally provides a function `shm_event_reconstruction` which decodes the SHM strings into a list of dataframes with the column names “location”, “pre_SHM_nt” and “post_SHM_nt”.

### immuneSIM random

The random mode allows for the simulation of fully random nucleotide and amino acid sequences of a specified length distribution. These sequences do not take into account V, D, J germline gene sequences and insertion or deletion information and are thus VDJ germline gene agnostic and can serve as a simple negative control. The output repertoire dataframe thus only contains information in the sequence and clone count columns of the repertoire dataframe while the remaining columns are set to `NA`.

### Modification of V,(D),J usage

immuneSIM provides three ways of introducing modified V, D, J frequencies into repertoires. First, through the modification of the reference list `list_germline_genes_allele_01` used for the germline gene sampling. A second option is provided by the `vdj_dropout` parameter of the immuneSIM function lets the user drop a chosen number of V, D and J genes (i.e., by setting their frequencies to zero). Finally, the user can also introduce noise into the VDJ frequencies via the parameter `vdj_noise` in the immuneSIM function. The noise value may be chosen between 0 (no noise) and 1 (recommended maximum noise) and sets the standard deviation for a normal distribution (mean = 0, bounded by −1,1) from which noise terms are sampled. Each sampled noise term is scaled by the frequency of the current germline gene and added to it (multiplicative noise). The `vdj_noise`, therefore, leads to a smaller modification of the germline gene frequencies than the first two options.

### Simulation of clonal abundance

Clonal abundance, shown to be a static representation of the clonal dynamics of immune repertoires that follows a power-law distribution (Greiff *et al.*, 2015; Greiff, Menzel, *et al.*, 2017), is simulated using the poweRlaw R package (Gillespie, 2015). Subsequently, the distribution of frequencies is transformed into counts by setting the lowest frequency to count = 1 (singletons). Unlike the simulated sequences, these counts are not based on experimental data. Therefore, there is no connection between a given sequence and its count as it is the case for the synthetic data provided by IGoR (Marcou *et al.*, 2017). The user can set the alpha parameter that controls the evenness of the power-law distribution and also has the choice to create a clone abundance that is equal across all clones (uniform distribution).

### Paired repertoires

The simulation of paired repertoires (heavy and light chain B-cell receptor repertoires, beta and alpha chain T-cell receptor repertoires) is achieved by simulating repertoires for heavy and light (or beta and alpha) receptor chains separately and combining them post-simulation using the immuneSIM function `combine_into_paired` which combines the repertoires and renames the columns. As the clonal frequencies are simulated in a sequence agnostic manner the heavy chain frequencies are kept while the frequencies from the light chain simulation discarded. The pairing occurs randomly as there is as of today only limited reliable information available on the preferential associations of chain pairing (Shcherbinin *et al.*, 2019; Zhou and Kleinstein, 2019).

### Simulation of standard and aberrant repertoires

The repertoires described in Supplementary Figures 2–5 contain 10’000 simulated sequences each. For the standard repertoires, default parameters of immuneSIM (representing experimental germline gene frequencies, no somatic hypermutation, no restrictions on insertion and deletions) were used. For the aberrant repertoires V, D, J usage was modified by introducing noise (`vdj_noise` = 0.2) and `vdj_dropout` depending on available germline genes (mm_igh: V = 10, D =3, J = 1, hs_igh: V = 10, D = 4, J = 2, mm_trb: V= 10, D = 1, J = 5 and hs_trb: V = 10, D = 0, J = 2). Additionally, insertions and deletions were restricted by disallowing n1 insertions (`ins_del_dropout =“no_insertions_n1”`). Finally, for IgH repertoires SHM was introduced (`shm.mode =“data”, shm.prob =15/350`) (See also Supplementary Table 2).

### Determining congruence with IMGT annotation

The full nucleotide VDJ sequences of each simulated repertoire were uploaded to and annotated with IMGT High-VQuest (Aouinti *et al.*, 2015). Following annotation, the percentage of exactly recovered immuneSIM V, D and J calls and the insertions and deletions by High-VQuest was determined. Further, the percentage of simulated sequences considered as in-frame and productive by IMGT as well as the substring and full string overlap of CDR3 sequences was calculated.

### Statistical analysis and plots

Statistical analysis was performed using R 3.4.0 (R Core Team). Graphics were generated using the ggplot2 (Wickham, 2009), ggthemes (Arnold, 2019), ComplexHeatmap (Gu *et al.*, 2016), circlize (Gu *et al.*, 2014) and RColorBrewer (Neuwirth, 2014) R packages and the ggplot2 theme theme.akbar (Akbar, 2019). The Supplementary Figures 2–5 were generated using the immuneSIM functions `plot_repertoire_A_vs_B and plot_report_repertoire`.

### Positional amino acid distribution

The amino acid frequencies were determined per position across all sequences of each CDR3 length. For the comparison of amino acid frequency distributions, the mean of the mean squared error (mmse) between distributions was calculated per position as previously described (Mason *et al.*, 2018). For the comparison between simulated and experimental repertoires, the CDR3s as annotated by IMGT High-VQuest (Aouinti *et al.*, 2015) were used.

### Gapped k-mer occurrence

For each repertoire, the occurrence of gapped k-mers was calculated across all CDR3 nucleotide sequences for parameters (k = 3, m ≤ 3, where k is the k-mer amino acid length and m is the number of amino acid gaps), as described by Palme *et al.*, 2015. The gapped-kmers are counted for the core of the CDR3 only, excluding the first three and last two amino acids containing more conserved patterns. The gapped k-mer occurrences were compared between simulated and experimental reference datasets (see Supplementary Table 1) and the Pearson and Spearman correlation was calculated. For the comparison between simulated and experimental repertoires, the CDR3s as annotated by IMGT High-VQuest (Aouinti *et al.*, 2015) were used.

### V, D, J germline gene usage

The usage of IMGT germline genes was compared between simulated and experimental reference datasets (see Supplementary Tables 1 and 2) and the Pearson and Spearman correlation coefficient was calculated. For this comparison, the V, D, J germline gene annotation from the immuneSIM output was used.

### Runtime and repertoire overlap

In order to evaluate the runtime performance of immuneSIM, five repertoires were generated using default parameters for each category (murine igh, murine trb, human igh, human trb) and repertoire size (10, 100, 1 ‘000, 10’000, 100’000 sequences). The runtime for the generation of each repertoire was measured using the tictoc R package (Izrailev, 2019) in R version 3.5 (R Core Team) on an iMac (Late 2013, Processor: 2.9 GHz Intel Core i5, Memory: 16GB 1600 MHz DDR3, macOS Sierra v 10.12.6). Subsequently, the repertoire overlap was calculated for each category across the five simulated repertoires as previously described (Greiff, Weber, *et al.*, 2017).

### Reference data

Germline gene sequences for T-cell receptor beta and alpha chains and B-cell receptor heavy and light chains were obtained from the IMGT reference directory (Alamyar *et al.*, 2012) on July 31, 2019 and collected in a reference R list. Subsequently, V, D, J usage frequencies from published data were added for human and murine T-cell receptor beta and B-cell receptor heavy chain germline genes while uniform distributions were added to human and murine T-cell receptor alpha and B-cell receptor light chain germline genes (see Supplementary Table 1, below).

**Supplementary Table. 1.**
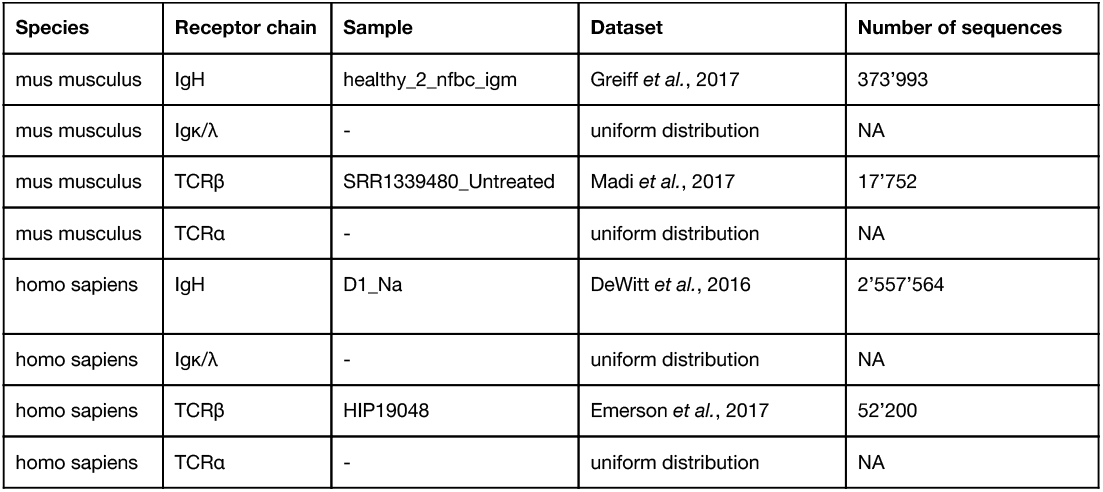
ImmuneSIM provides germline gene frequencies based on four published datasets. For the simulation of T-cell receptor alpha chains and B-cell receptor light chains, immuneSIM utilizes uniform sampling of IMGT germline genes since the number of available datasets is currently too limited to compute reliable germline frequencies.

**Supplementary Table. 2.** Parameters for the simulation of standard and aberrant repertoires. Standard repertoires were simulated using default parameters based on the experimental reference datasets. For the aberrant repertoires, noise was introduced into the VDJ germline frequencies, VDJ germline genes were dropped out, n1 insertions were disallowed and for aberrant IgH repertoires SHM was introduced.

| Supplementary Table 2. Parameters for standard and aberrant repertoires. (SFig. 2—7) |  |  |  |  |  |  |
| --- | --- | --- | --- | --- | --- | --- |
| Name | species, receptor, chain | vdj_noise | vdj_dropout | ins_del_dropout | shm.mode | shm.prob |
| standard murine igh | "mm", "ig", "h" | 0 | V = 0, D = 0, J = 0 | "" | "none" | - |
| aberrant murine igh | "mm", "ig", "h" | 0.2 | V = 10, D = 3, J = 1 | "no_insertions_n1" | "data" | 15/350 |
| standard murine trb | "mm", "tr", "b" | 0 | V = 0, D = 0, J = 0 | "" | "none" | - |
| aberrant murine trb | "mm", "tr", "b" | 0.2 | V = 10, D = 1, J = 5 | "no_insertions_n1" | "none" | - |
| standard human igh | "hs", "ig", "h" | 0 | V = 0, D = 0, J = 0 | "" | "none" | - |
| aberrant human igh | "hs", "ig", "h" | 0.2 | V = 10, D = 4, J = 2 | "no_insertions_n1" | "data" | 15/350 |
| standard human trb | "hs", "tr", "b" | 0 | V = 0, D = 0, J = 0 | "" | "none" | - |
| aberrant human trb | "hs", "tr", "b" | 0.2 | V = 10, D = 0, J = 2 | "no_insertions_n1" | "none" | - |

---

**SFig. 1.**
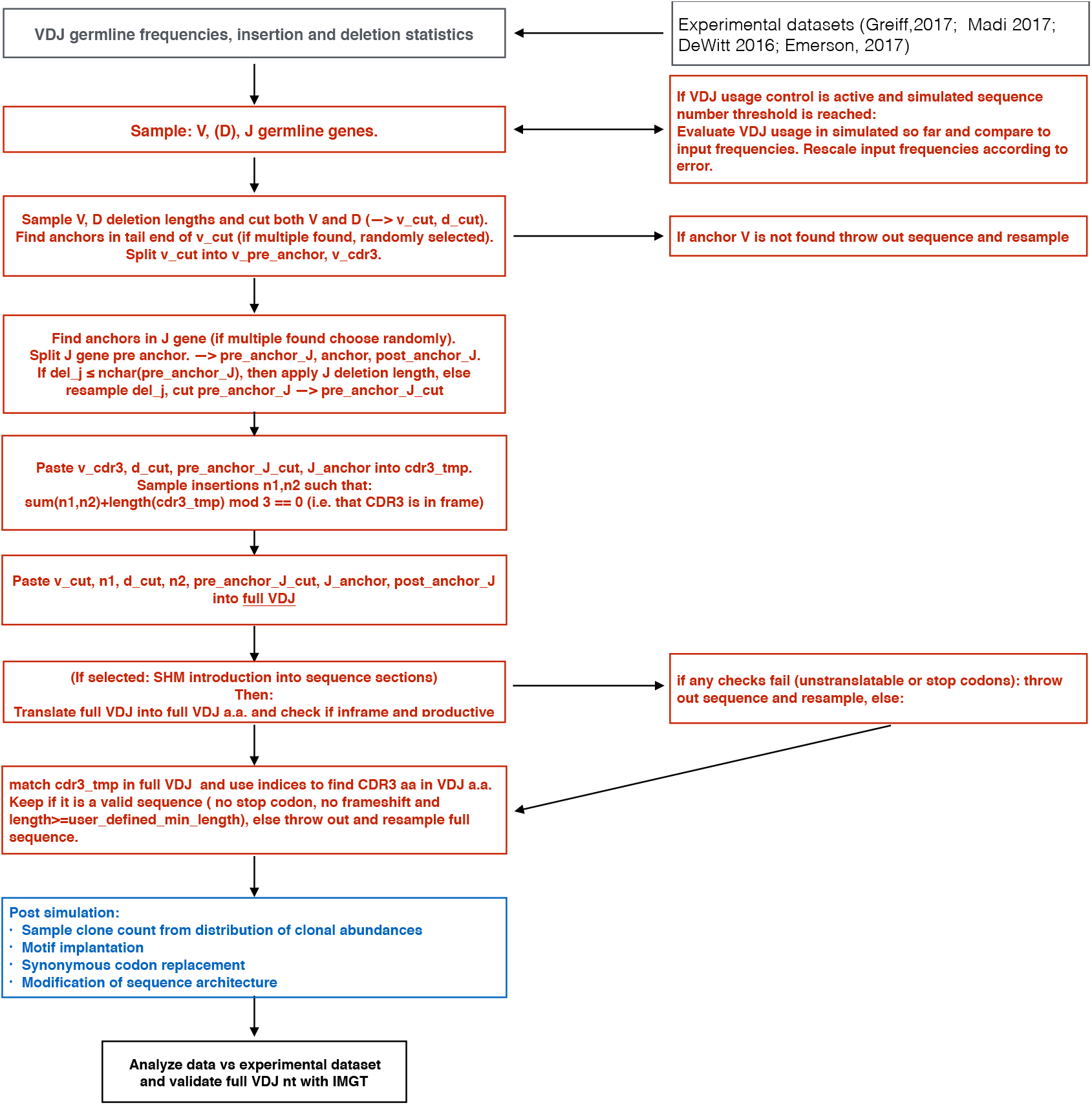
Flowchart of immune receptor simulation in the R package immuneSIM. immuneSIM allows simulation of immune repertoires via an in silico V(D)J recombination algorithm with input parameters (VDJ frequencies, insertion sequences and deletion lengths) in a user-defined fashion based on experimental data. The germline V, D and J genes are sampled according to usage frequencies defined in the input. After germline gene sampling, a CDR3 anchor is identified in the V and J gene. Subsequently, V and D and J deletion lengths are sampled randomly. Finally, the n1 and n2 insertions are sampled from a subset of insertions such that their length complements the current preliminary CDR3 resulting in an in-frame CDR3. The resulting nucleotide sequence is subsequently translated and either discarded (if it contains a stop codon) or kept as a valid in silico sequence. In the case of B-cell receptor simulations, SHM are introduced based on the user-defined parameters.

**SFig. 2.**
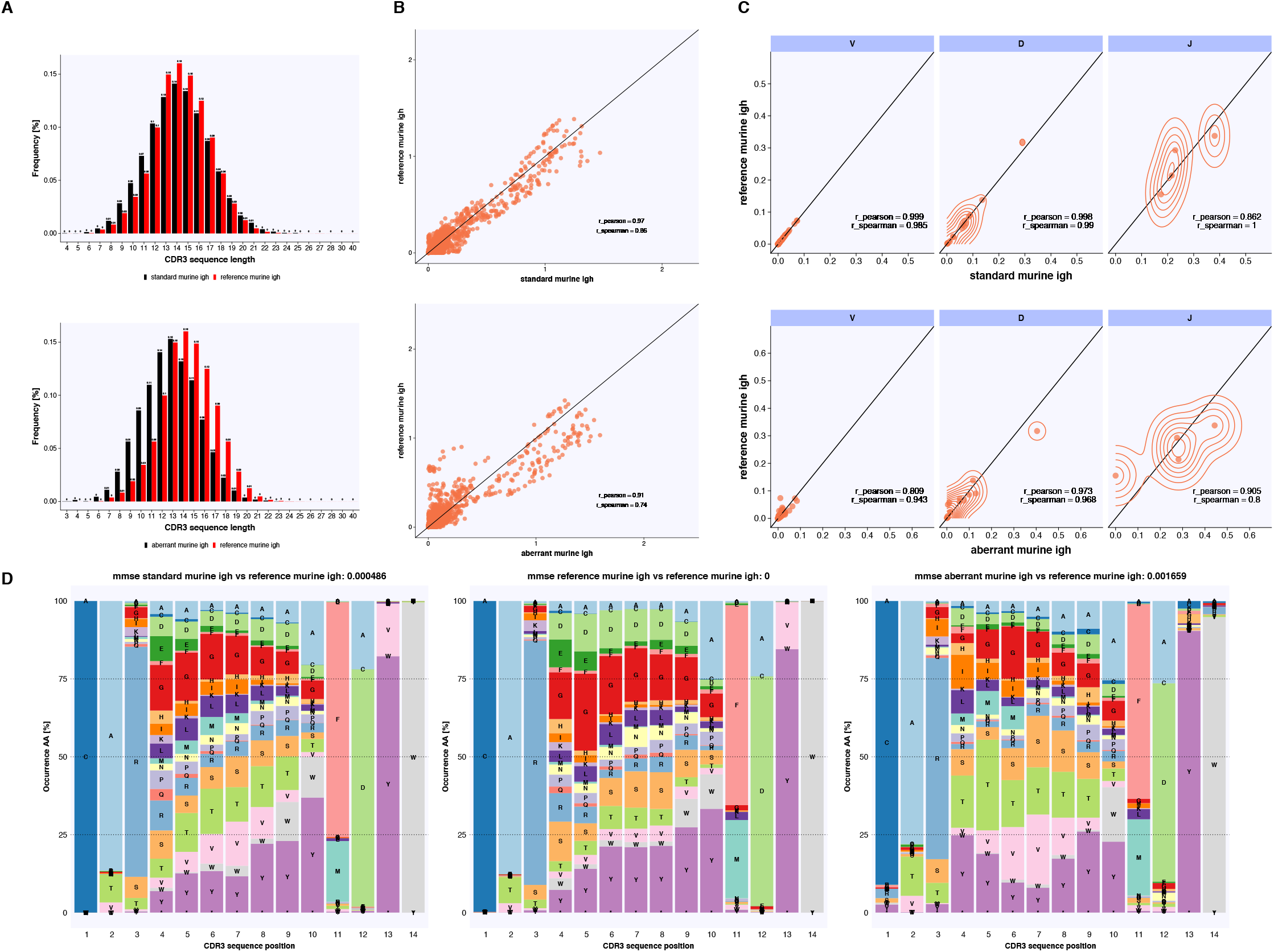
immuneSIM simulates fully annotated murine IgH immune repertoires that are either high similar or deviating from experimental immune repertoires in regards to the major immune repertoire features length distribution, gapped-kmer occurrence, VDJ usage and CDR3 amino acid frequency. (A) The CDR3 length distributions of a standard simulated murine IgH repertoire (simulated using default parameters) and experimental data (both annotated using IMGT) largely overlap, while the aberrant repertoire simulated with non-default parameters (Supplementary Table 2) shows a shift towards shorter lengths. (B) Gapped k-mer occurrence of CDR3 nucleotide sequence shows high correlation between default parameter simulation and experimental repertoires (upper panel, r_spearman_ = 0.86 for k = 3 and gap size m ≤ 3, nkmers = 16384) and lower correlation to aberrant repertoires (lower panel, r_spearman_ = 0.74). (C) The V, D and J frequencies between simulated (default parameters) and experimental repertoires are highly correlated (r_spearman_ ≥ 0.985). Simulating aberrant repertoires is also possible (lower panel, r_spearman_ ≥ 0.8). (D) The positional amino acid frequencies of CDR3 sequences (annotated using IMGT) of length 14 are shown to be highly similar (Mean of mean squared errors across positions, mmse: 0.000486) to experimental data (center). Repertoires simulated to be farther from the experimental dataset deviating with respect to positional amino acid frequency (mmse: 0.001659).

**SFig. 3.**
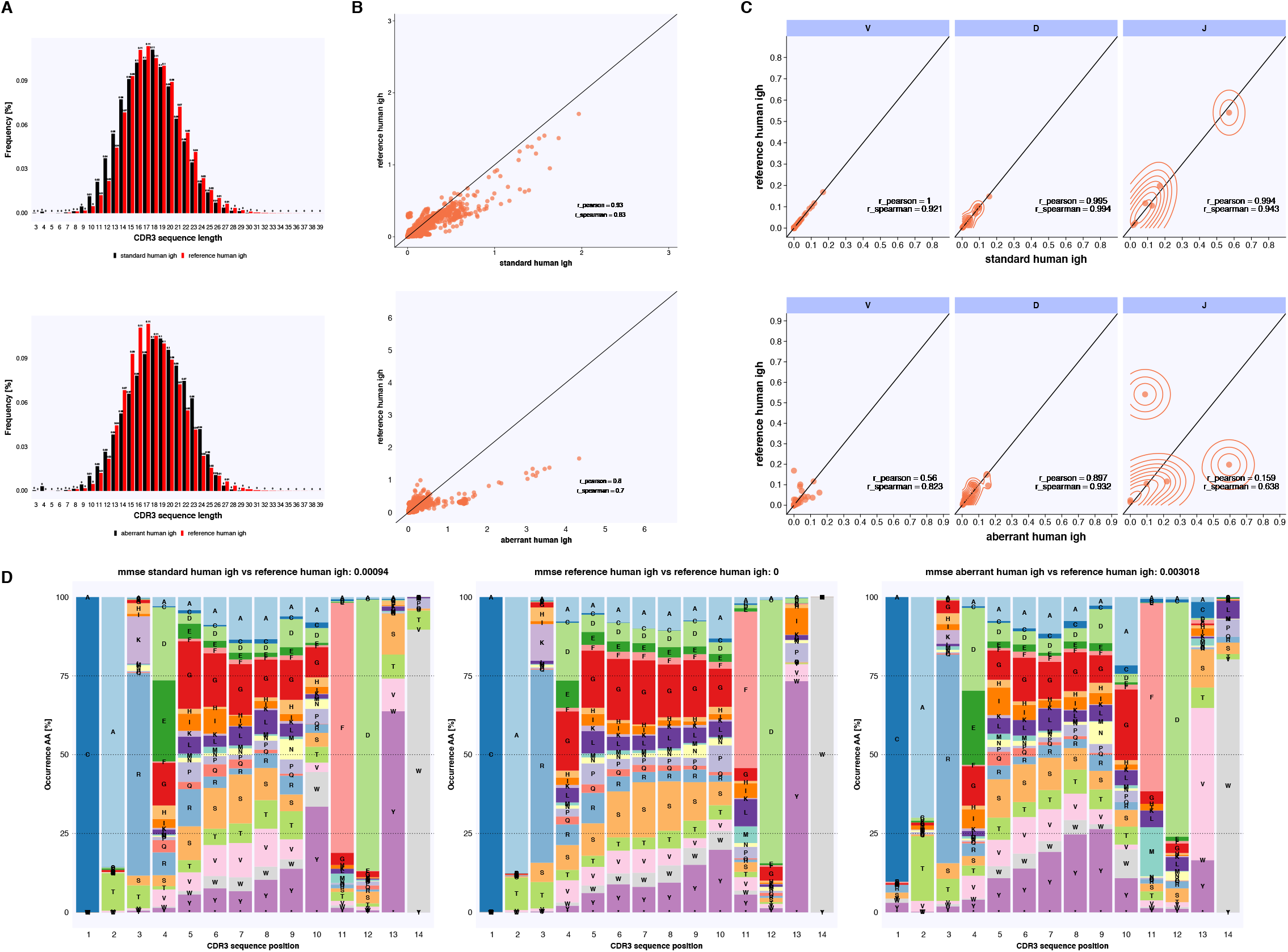
Major features of standard and aberrant simulated human IgH repertoires. (A) The CDR3 length distributions of a standard simulated repertoires (simulated using default parameters) and experimental data (both annotated using IMGT) largely overlap while the aberrant repertoire simulated with non-default parameters (Supplementary Table 2) shows a shift towards shorter lengths. (B) Gapped k-mer occurrence of CDR3 nucleotide sequence shows high correlation between simulated (default parameters) and experimental repertoires (upper panel, r_spearman_ = 0. 83 for k = 3 and gap size m ≤ 3, nkmers = 16384) and lower correlation to aberrant repertoires (lower panel, r_spearman_ = 0.7). (C) The V, D and J frequencies between simulated (default parameters) and experimental repertoires correlate to a high degree (r_spearman_ ≥ 0.921). Simulating more deviating repertoires is also possible (lower panel, r_spearman_ ≥ 0.638). (D) The positional amino acid frequencies of CDR3 sequences (annotated using IMGT) of length 14 are shown to be highly similar (Mean of mean squared errors across positions, mmse: 0.00094) to experimental data (center). Repertoires simulated to be farther from the experimental dataset deviating with respect to positional amino acid frequency (mmse: 0.003018).

**SFig. 4.**
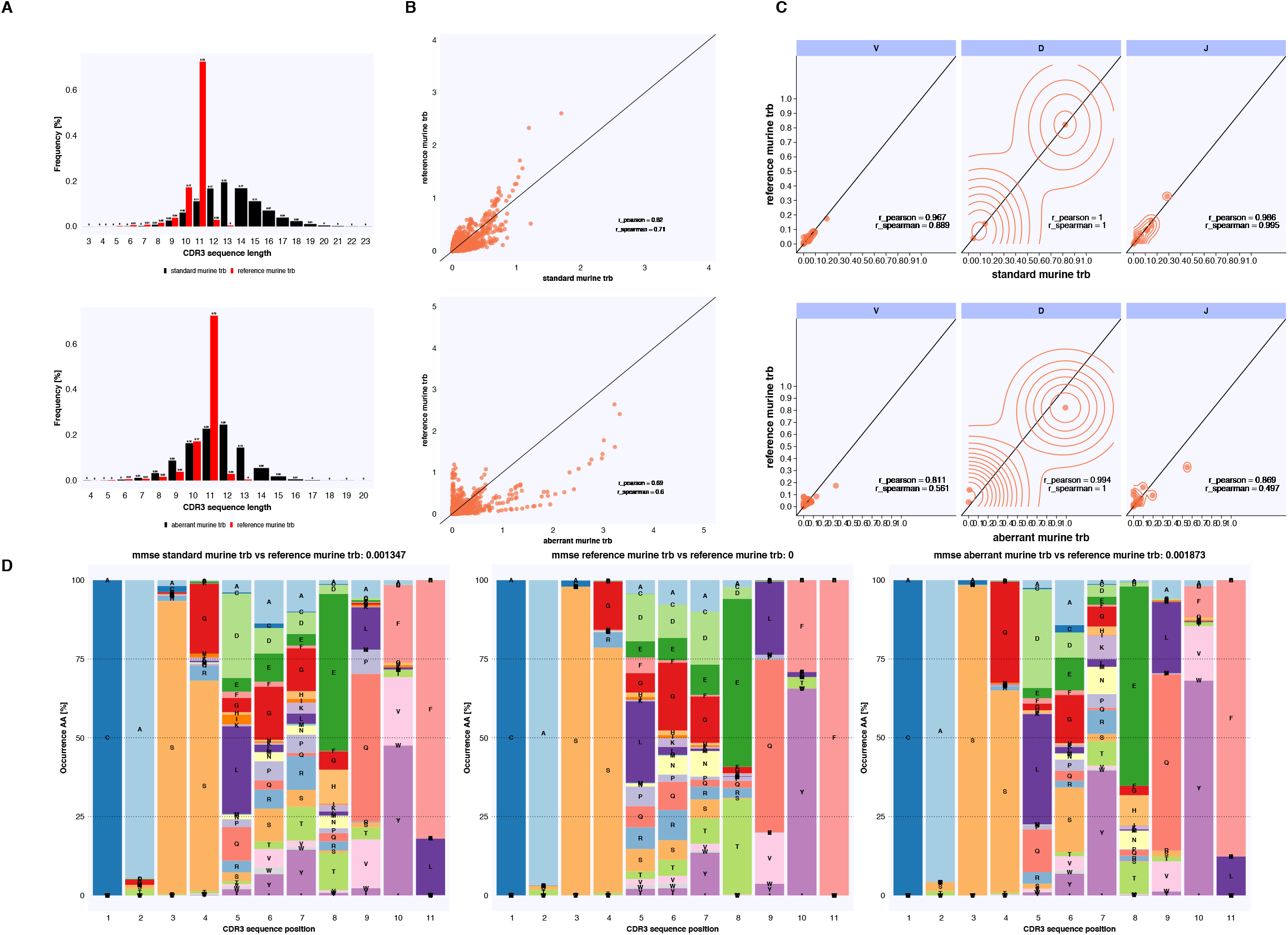
Major features of standard and aberrant simulated murine TCRβ repertoires. The recovery of murine trb features are more difficult using immuneSIM due to the generally shorter sequence length. Nevertheless standard simulated repertoires (simulated using default parameters) and a aberrant repertoires simulated with non-default parameters (Supplementary Table 2) can be generated with different levels of similarity to experimental data. (A) The CDR3 length distributions of repertoires simulated with default parameters and experimental data (both annotated using IMGT) are shifted with the simulated repertoire having a more even distribution. (B) Gapped k-mer occurrence of CDR3 nucleotide sequence shows higher correlation between default parameter simulation and experimental repertoires (upper panel, r_spearman_ = 0.71 for k = 3, m ≤ 3, nkmers = 16384) and low correlation to aberrant repertoires (lower panel, r_spearman_ = 0.6). (C) The V, D and J frequencies between simulated (default parameters) and experimental repertoires correlate to a high degree (upper panel, r_spearman_ ≥ 0.889). Simulating more deviating repertoires is also possible (lower panel, r_spearman_ ≥ 0.497). (D) The positional amino acid frequencies of CDR3 sequences (annotated using IMGT) of length 14 are shown to be highly similar (left, Mean of mean squared errors across positions, mmse: 0.001347) to experimental data (center). Repertoires simulated to be farther from the experimental dataset deviating with respect to positional amino acid frequency (right, mmse: 0.001873).

**SFig. 5.**
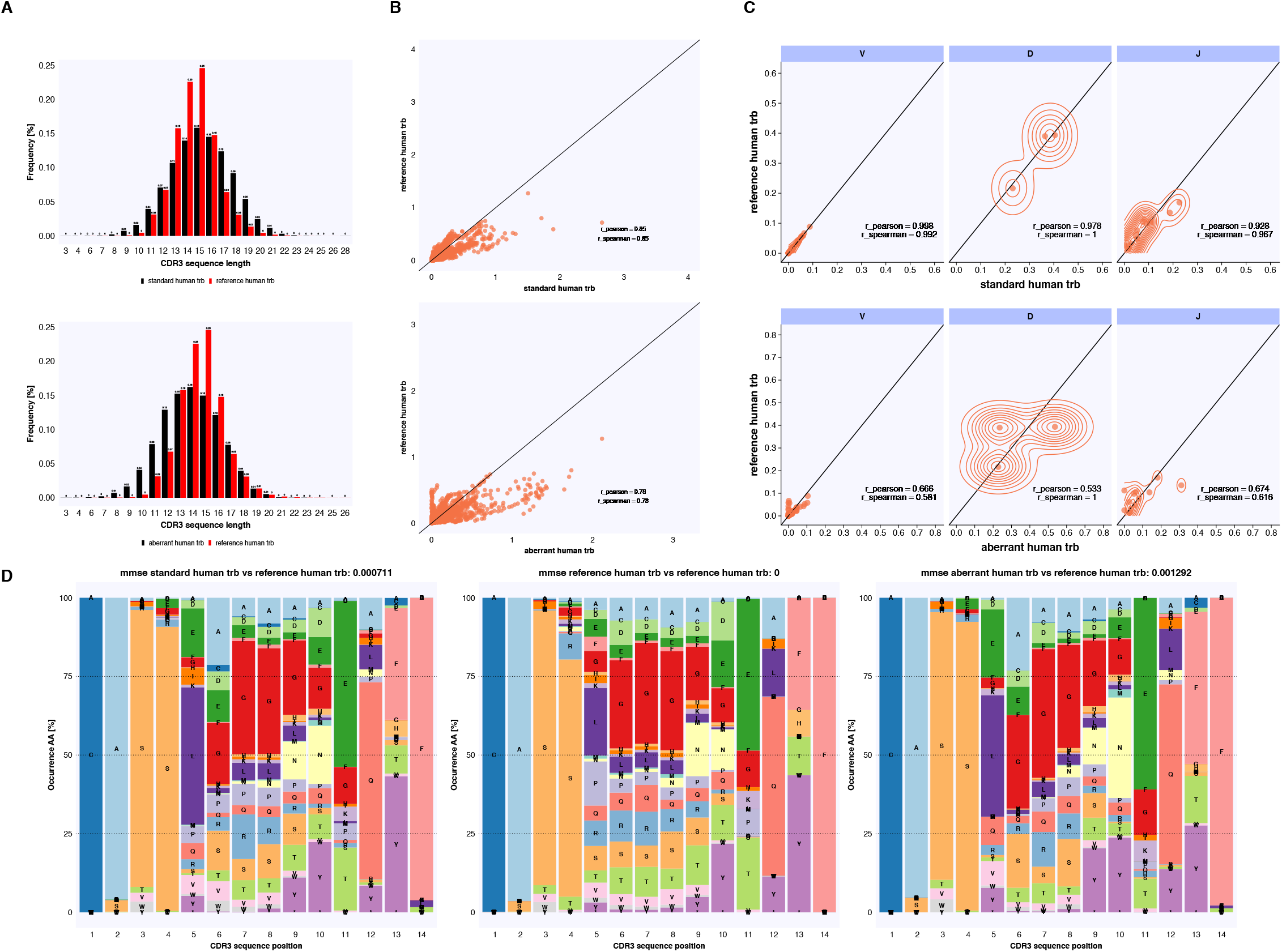
Major features of standard and aberrant simulated human TCRβ repertoires. (A) The CDR3 length distributions of a standard repertoire (simulated using default parameters) and experimental data (both annotated using IMGT) largely overlap while the aberrant repertoire simulated with non-default parameters (Supplementary Table 2) shows a shift to shorter lengths. (B) Gapped k-mer occurrence of CDR3 nucleotide sequence shows high correlation between simulated (standard) and experimental repertoires (upper panel, r_spearman_ = 0.85 for k = 3, m ≤ 3, nkmers = 16384) and lower correlation to aberrant repertoires (lower panel, r_spearman_ = 0.78). (C) The V, D and J frequencies between standard simulated (default parameters) and experimental repertoires correlate to a high degree (r_spearman_ ≥ 0.967). Simulating more deviating repertoires is also possible (lower panel, r_spearman_ ≥ 0.581). (D) The positional amino acid frequencies of CDR3 sequences (annotated using IMGT) of length 14 are shown to be highly similar (Mean of mean squared errors across positions, mmse: 0.000711) to experimental data (center). Repertoires simulated to be farther from the experimental dataset deviating with respect to positional amino acid frequency (mmse: 0.001292).

**SFig. 6.**
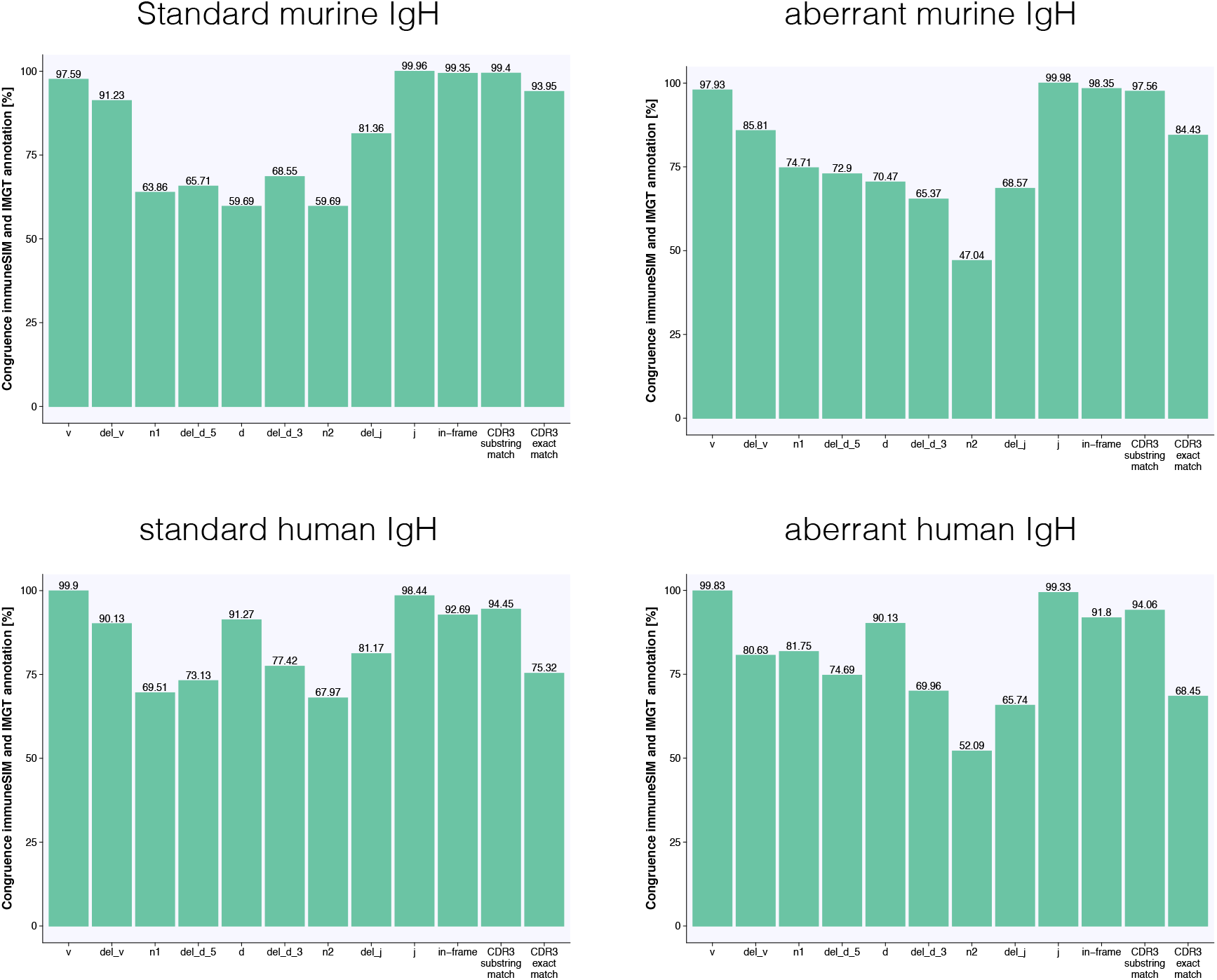
ImmuneSIM generates B-cell receptor sequences that are in accord with IMGT annotation. Simulation of murine (top) and human (bottom) IgH repertoires each using default (standard, left) and nondefault parameters (aberrant, right). The comparison of immuneSIM-annotated simulated repertoires with IMGT HighV-Quest annotation indicates that immuneSIM generates productive sequences with IMGT identifiable V,D,J genes, insertions, and deletions. The lower percent of congruence with respect to the n1-D-n2 portion of the CDR3 is expected due to the difficulty of D-gene annotation (Bolotin *et al.*, 2015). Nearly all immuneSIM-simulated immune receptor sequences have an identifiable in-frame (>91.8%) CDR3 junction that is correctly annotated by immuneSIM to the degree that it is equal to or a substring of the IMGT CDR3 in >94.06% of all cases. N = 10’000.

**SFig. 7.**
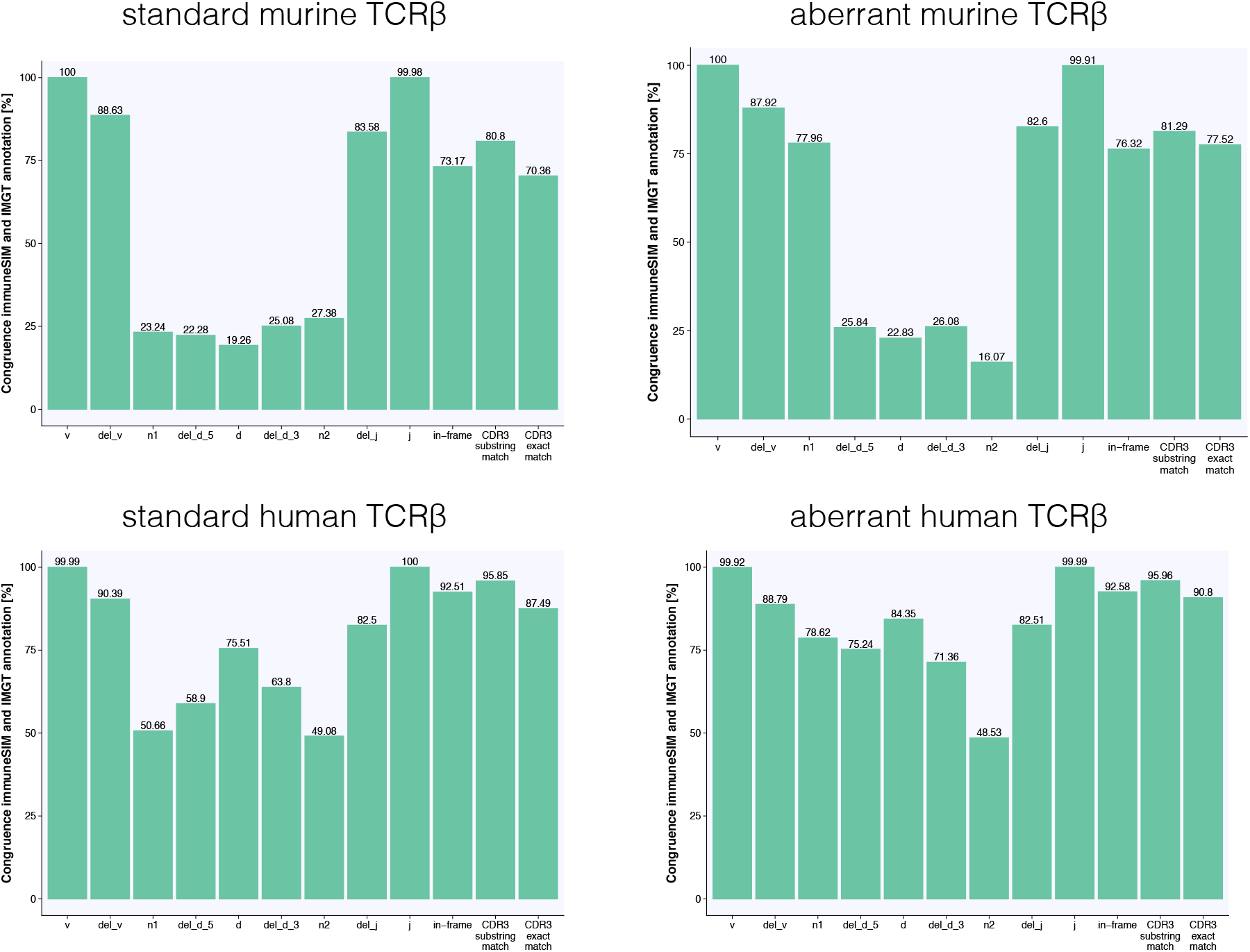
High congruence with IMGT annotation is high for TCRβ repertoires across multiple species. Simulation of murine (top) and human (bottom) TCRβ repertoires each using default (standard, left) and nondefault (aberrant, right) parameters. The comparison of immuneSIM-annotated simulated repertoires with IMGT HighV-Quest annotation indicates that immuneSIM generates productive sequences that can be annotated by IMGT. The performance is lowest for murine TCRβ, likely due to the increased difficulty the shorter CDR3s present especially with regard to D-gene annotation. However, immuneSIM repertoires have recognizable V and J genes (~100%) and >73% of sequences are determined to be in-frame with >70% exact CDR3 matches between IMGT and immuneSIM.

**SFig. 8.**
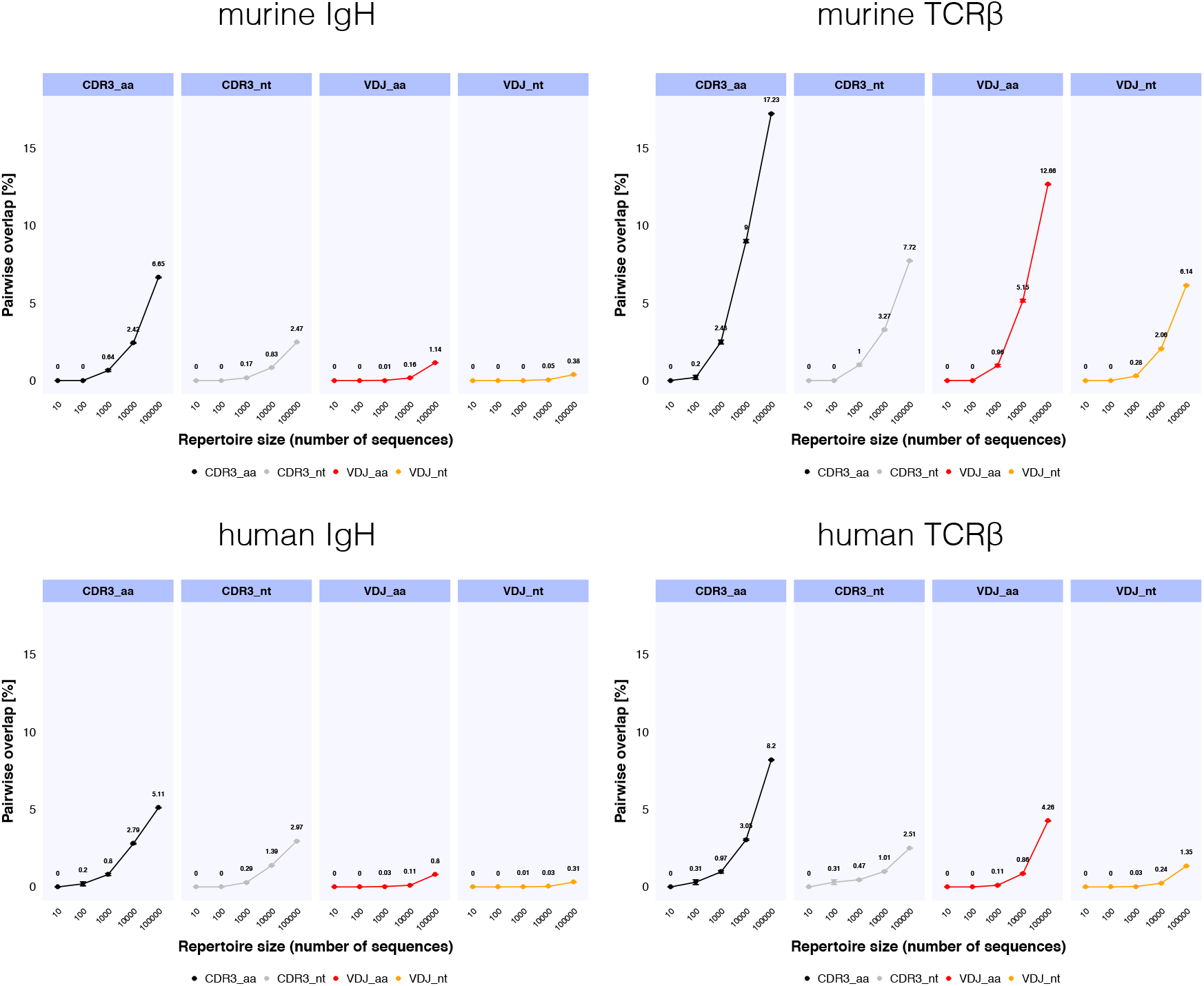
The pairwise sequence overlap across simulated repertoires increases with repertoire size. Pairwise overlap between five repertoires per repertoire size and species/receptor chain combination was measured for amino acid and nucleotide CDR3 and VDJ sequences. Overlap reaches a maximum of 17.23% for the repertoire size of 100’000 murine TCRβ CDR3 amino acid sequences, confirming that immuneSIM produces highly diverse immune receptor repertoire sequences.

**SFig.9.**
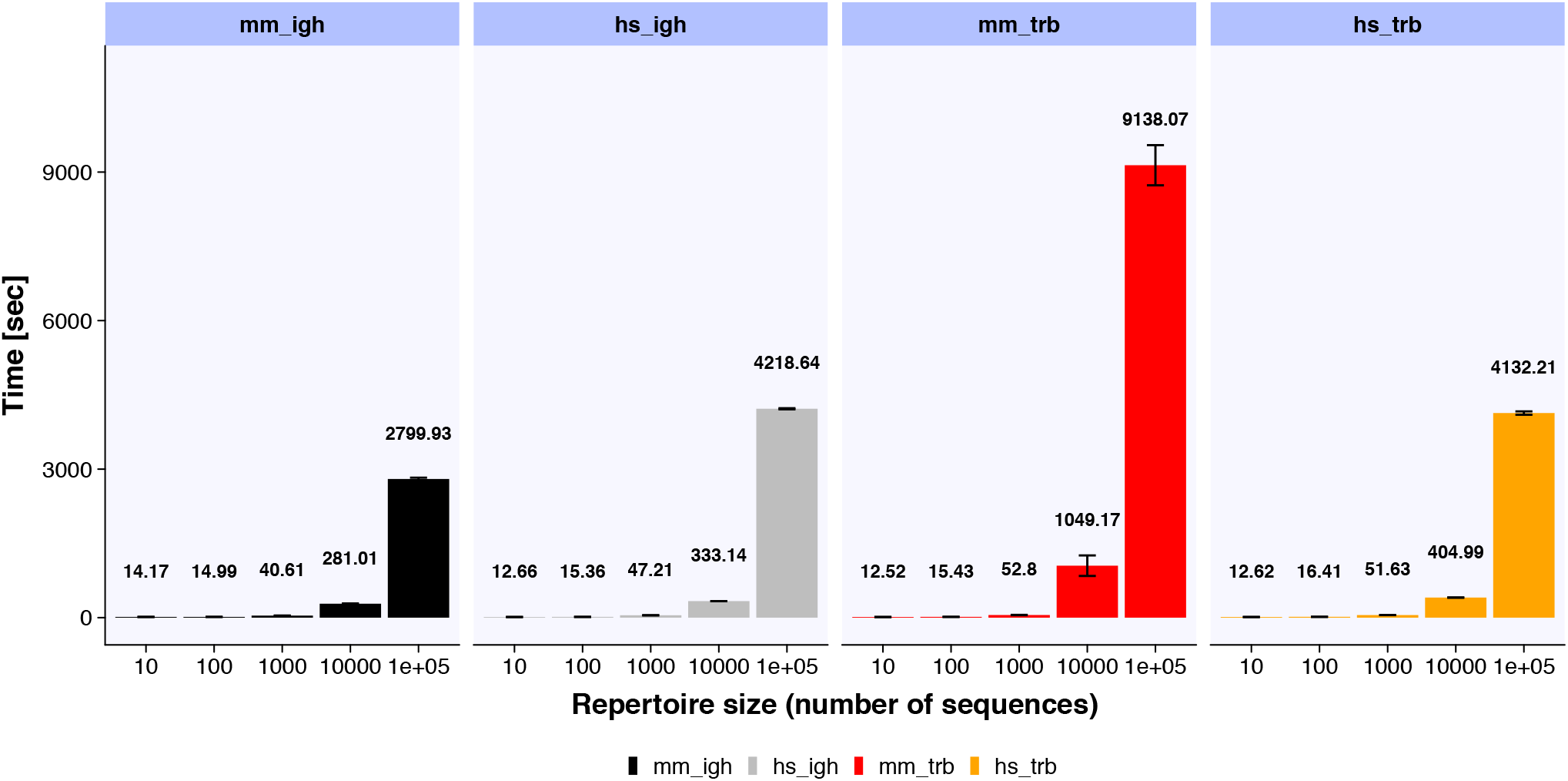
immuneSIM runtime differs for various species/receptor chain combinations. Murine and human IgH repertoires (mm_igh, hs_igh) can be simulated efficiently while in silico recombination of murine and human TCRβ repertoires (mm_trb, hs_trb) requires more time for larger datasets (due to lower likelihood of in-frame, productive recombination events). Runtime was measured as an average across five standard parameter simulations per size and category using the tictoc R package.

## References

Alamyar,E. et al. (2012) IMGT® Tools for the Nucleotide Analysis of Immunoglobulin (IG) and T Cell Receptor (TR) V-(D)-J Repertoires, Polymorphisms, and IG Mutations: IMGT/V-QUEST and IMGT/HighV-QUEST for NGS. In, Christiansen,F.T. and Tait,B.D. (eds), Immunogenetics, Methods and Applications in Clinical Practice, Methods in Molecular Biology. Humana Press, Totowa, NJ, pp. 569–604.

Arora,R. et al. (2019) Repertoire-Based Diagnostics Using Statistical Biophysics. bioRxiv, 519108.

Bashford-Rogers,R.J. et al. (2013) Network properties derived from deep sequencing of human B-cell receptor repertoires delineate B-cell populations. Genome Res., 23, 1874–1884.

Bolotin,D.A. et al. (2015) MiXCR: software for comprehensive adaptive immunity profiling. Nat. Methods, 12, 380–381.

Breden,F. et al. (2017) Reproducibility and Reuse of Adaptive Immune Receptor Repertoire Data. Front. Immunol., 8.

Brown,A.J. et al. (2019) Augmenting adaptive immunity: progress and challenges in the quantitative engineering and analysis of adaptive immune receptor repertoires. Mol. Syst. Des. Eng., 4, 701–736.

Dash,P. et al. (2017) Quantifiable predictive features define epitope-specific T cell receptor repertoires. Nature, 547, 89–93.

DeWitt,W.S. et al. (2016) A Public Database of Memory and Naive B-Cell Receptor Sequences. PLOS ONE, 11, e0160853.

Emerson,R.O. et al. (2017) Immunosequencing identifies signatures of cytomegalovirus exposure history and HLA-mediated effects on the T cell repertoire. Nat. Genet., 49, 659–665.

Giudicelli,V. et al. (2004) IMGT/V-QUEST, an integrated software program for immunoglobulin and T cell receptor V-J and V-D-J rearrangement analysis. Nucleic Acids Res., 32, W435–W440.

Giudicelli,V. and Lefranc,M.-P. (2011) IMGT/JunctionAnalysis: IMGT Standardized Analysis of the V-J and V-D-J Junctions of the Rearranged Immunoglobulins (IG) and T Cell Receptors (TR). Cold Spring Harb. Protoc., 2011, pdb.prot5634-pdb.prot5634.

Glanville,J. et al. (2017) Identifying specificity groups in the T cell receptor repertoire. Nature, 547, 94–98.

Greiff,V. et al. (2015) Bioinformatic and Statistical Analysis of Adaptive Immune Repertoires. Trends Immunol., 36, 738–749.

Greiff,V., Weber,C.R., et al. (2017) Learning the High-Dimensional Immunogenomic Features That Predict Public and Private Antibody Repertoires. J. Immunol. Baltim. Md 1950, 199, 2985–2997.

Greiff,V., Menzel,U., et al. (2017) Systems Analysis Reveals High Genetic and Antigen-Driven Predetermination of Antibody Repertoires throughout B Cell Development. Cell Rep., 19, 1467–1478.

Madi,A. et al. (2017) T cell receptor repertoires of mice and humans are clustered in similarity networks around conserved public CDR3 sequences. eLife, 6, e22057.

Madi,A. et al. (2014) T-cell receptor repertoires share a restricted set of public and abundant CDR3 sequences that are associated with self-related immunity. Genome Res., 24, 1603–1612.

Marcou,Q. et al. (2017) IGoR: A Tool For High-Throughput Immune Repertoire Analysis. bioRxiv, 141143.

Miho,E. et al. (2018) Computational strategies for dissecting the high-dimensional complexity of adaptive immune repertoires. Front. Immunol., 9.

Miho,E. et al. (2017) The fundamental principles of antibody repertoire architecture revealed by large-scale network analysis. bioRxiv, 124578.

Olson,B.J. et al. (2019) sumrep: a summary statistic framework for immune receptor repertoire comparison and model validation. bioRxiv, 727784.

Ostmeyer,J. et al. (2019) Biophysicochemical Motifs in T-cell Receptor Sequences Distinguish Repertoires from Tumor-Infiltrating Lymphocyte and Adjacent Healthy Tissue. Cancer Res., 79, 1671–1680.

Safonova,Y. et al. (2015) IgSimulator: a versatile immunosequencing simulator. Bioinformatics, btv326.

Safonova,Y. and Pevzner,P.A. (2019) De novo inference of diversity genes and analysis of non-canonical V(DD)J recombination in immunoglobulins. ArXiv 190102483 Q-Bio.

Yaari,G. and Kleinstein,S.H. (2015) Practical guidelines for B-cell receptor repertoire sequencing analysis. Genome Med., 7, 121.

Yermanos,A. et al. (2017) Comparison of methods for phylogenetic B-cell lineage inference using time-resolved antibody repertoire simulations (AbSim). Bioinformatics.

## References

Akbar,R. (2019) themeakbar Zenodo.

Alamyar,E. et al. (2012) IMGT® Tools for the Nucleotide Analysis of Immunoglobulin (IG) and T Cell Receptor (TR) V-(D)-J Repertoires, Polymorphisms, and IG Mutations: IMGT/V-QUEST and IMGT/HighV-QUEST for nGs. In, Christiansen,F.T. and Tait,B.D. (eds), Immunogenetics: Methods and Applications in Clinical Practice, Methods in Molecular Biology. Humana Press, Totowa, NJ, pp. 569–604.

Aouinti,S. et al. (2015) IMGT/HighV-QUEST Statistical Significance of IMGT Clonotype (AA) Diversity per Gene for Standardized Comparisons of Next Generation Sequencing Immunoprofiles of Immunoglobulins and T Cell Receptors. PLoS ONE, 10

Arnold,J.B. (2019) ggthemes: Extra Themes, Scales and Geoms for ‘ggplot2’.

Csardi,G. and Nepusz,T. (2006) The igraph software package for complex network research. InterJournal, Complex Systems

Emerson,R.O. et al. (2017) Immunosequencing identifies signatures of cytomegalovirus exposure history and HLA-mediated effects on the T cell repertoire. Nat. Genet., 49

Gillespie,C.S. (2015) Fitting Heavy Tailed Distributions: The poweRlaw Package. J. Stat. Softw., 64

Glanville,J. et al. (2017) Identifying specificity groups in the T cell receptor repertoire. Nature, 547

Greiff,V. et al. (2015) A bioinformatic framework for immune repertoire diversity profiling enables detection of immunological status. Genome Med., 7

Greiff,V., Weber,C.R., et al. (2017) Learning the High-Dimensional Immunogenomic Features That Predict Public and Private Antibody Repertoires. J. Immunol. Baltim. Md 1950, 199

Greiff,V., Menzel,U., et al. (2017) Systems Analysis Reveals High Genetic and Antigen-Driven Predetermination of Antibody Repertoires throughout B Cell Development. Cell Rep., 19

Gu,Z. et al. (2014) circlize Implements and enhances circular visualization in R. Bioinforma. Oxf. Engl., 30, 2811–2812.

Gu,Z. et al. (2016) Complex heatmaps reveal patterns and correlations in multidimensional genomic data. Bioinforma. Oxf. Engl., 32

Lefranc,M.-P. (2011) From IMGT-ONTOLOGY DESCRIPTION Axiom to IMGT Standardized Labels: For Immunoglobulin (IG) and T Cell Receptor (TR) Sequences and Structures. Cold Spring Harb. Protoc., 2011, pdb.ip83.

Loo,M.P.J. van der (2014) The stringdist Package for Approximate String Matching. R J., 6

Mason,D.M. et al. (2018) High-throughput antibody engineering in mammalian cells by CRISPR/Cas9-mediated homology-directed mutagenesis. Nucleic Acids Res.

Miho,E. et al. (2019) Large-scale network analysis reveals the sequence space architecture of antibody repertoires. Nat. Commun., 10

Neuwirth,E. (2014) RColorBrewer: ColorBrewer Palettes.

Ostmeyer,J. et al. (2019) Biophysicochemical Motifs in T-cell Receptor Sequences Distinguish Repertoires from Tumor-Infiltrating Lymphocyte and Adjacent Healthy Tissue. Cancer Res., 79

Palme,J. et al. (2015) KeBABS: an R package for kernel-based analysis of biological sequences: Fig. 1. Bioinformatics, 31

R Core Team R: A Language and Environment for Statistical Computing R Foundation for Statistical Computing, Vienna, Austria.

Rubelt,F. et al. (2017) Adaptive Immune Receptor Repertoire Community recommendations for sharing immune-repertoire sequencing data. Nat. Immunol.

Shcherbinin,D.S. et al. (2019) Comprehensive analysis of structural and sequencing data reveals almost unconstrained chain pairing in TCRαβ complex. bioRxiv, 693630.

Wickham,H. (2009) ggplot2: Elegant Graphics for Data Analysis Springer-Verlag New York.

Zhou,J.Q. and Kleinstein,S.H. (2019) Immunoglobulin heavy chains are sufficient to determine most B cell clonal relationships. bioRxiv, 665760.

